# Sugar shock: Probing *Streptococcus pyogenes* metabolism through bioluminescence imaging

**DOI:** 10.1101/2022.01.15.476453

**Authors:** Richard W. Davis, Charlotte G. Muse, Heather Eggleston, Micaila Hill, Peter Panizzi

**Affiliations:** Department of Drug Discovery and Development, Harrison School of Pharmacy, 720 South Donahue, Auburn University, Auburn, AL 36849

**Keywords:** bioluminescence, biosensors, pathogens, physiology, *Streptococcus pyogenes*, molecular imaging

## Abstract

*Streptococcus pyogenes* (*S. pyogenes*) can thrive in its host during an infection, and, as a result, it must be able to respond to external stimuli and available carbon sources. The preclinical use of engineered pathogens capable of constitutive light production may provide realtime information on microbial-specific metabolic processes. Here we mapped the central metabolism of a *luxABCDE*-modified *S. pyogenes* Xen20 (*Strep*. Xen20) to its *de novo* synthesis of luciferase substrates as assessed by the rate of light production in response to different environmental triggers. Previous characterization predicted that the *lux* operon was under the myo-inositol iolE promotor. Here we show that supplementation with myoinositol generated increased Xen20 luminescence. Surprisingly, when supplemented with infection-relevant carbon sources, such as glucose or glycine, light production was diminished. This was presumably due to the scavenging of pyruvate by *L*-lactate dehydrogenase (LDH). Inhibition of LDH by its inhibitor, oxamate, partially restored luminescent signal in the presence of glucose, presumably by allowing the resulting pyruvate to proceed to acetyl-coenzyme A (CoA). This phenomenon appeared specific to the lactic acid bacterial metabolism as glucose or glycine did not reduce signal in an analogous *luxABCDE*-modified Gram-positive pathogen, *Staph*. Xen29. The *Strep.* Xen20 cells produced light in a concentration-dependent manner, inversely related to the amount of glucose present. Taken together, our measures of microbial response could provide new information regarding the responsiveness of *S. pyogenes* metabolism to acute changes in its local environments and cellular health.

## Introduction

*Streptococcus pyogenes* (*S. pyogenes*) infections often manifest as necrotizing fasciitis or cellulitis (1). Surveillance Report for 2017 reported 23,650 new cases of *S. pyogenes* infections for every 33 million individuals. The incidence of cellulitis or necrotizing fasciitis over the same period was 44.5% and 4.9%, respectively (2). The total rate for 2017 showed 7.6 per 100,000 cases, and 8.4% of those resulted in death (2).

An intriguing pre-clinical method for real-time *in vivo* tracking of these harmful pathogens during an infection is bioluminescence. Although the use of engineered light-producing microbes has limited diagnostic value, the benefit of these types of bacteria to pre-clinical experimentation is often underappreciated. For example, deposition of bacterial-targeted dyes *in vivo* during pre-clinical testing can be readily verified by co-localization with the light signal from one of these engineered microbes. Typically, such studies require that the parental strain be transformed with plasmids containing either the *Photorhabdus luminescens lux* operon cassette (*luxABCDE*) or the firefly luciferase (*ffluc*) enzyme. One significant limitation to the use of these pathogens is the gap in our understanding regarding changes in light production by these microbes. The *luxABCDE* cassette often is placed on a transposable element, so expression is dependent on its random insertion into the genome of the parent bacterium (3). As such, light production can be linked to cellular processes that are inherent to the microbe and report on the microbes’ response to its local environments. Previously, we studied a *luxABCDE-*modified *Streptococcus pyogenes* Xen20 (*Strep*. Xen20) to track the spread of cutaneous infection in wild type mice (4). By *ex vivo* colony forming units (CFU) determination and Gram-staining histology, we noted live *Streptococcus* present in the distal organs of the infected animals. Our findings did not support the non-invasive bioluminescent imaging (BLI) results, suggesting either the signal was below the limit of detection or there was a breakdown of light production *in vivo*. We found that light production by *Strep.* Xen20 decreased with increasing *D*-glucose concentration; thereby, essentially serving as a glucose biosensor within the animal. Recently, Mimee *et al*. reported on a similar light-producing *Escherichia coli* (*E. coli*) used in an ingestible bacterial-electronic system that detects the presence of heme in a model of gastric bleeding (5). Such microbial systems or devices could serve as biosensors in venues ranging from clinical to industrial.

Light production by the emitting microbes follows the conversion of activated fatty acyl compounds to fatty aldehydes via the *luxCDE* system (6). These fatty aldehydes drive the *luxAB* complexes, which use molecular oxygen and reduced flavin mononucleotide (FMNH_2_) to produce fatty acid, water, oxidized flavin mononucleotide, and light. Therefore, an essential requirement of bioluminescence is the availability of critical reactants, such as nonanoic, FMNH_2_, and adenosine triphosphate (ATP) (7). Herein, we show a differential light production based on catabolite repression and gluconeogenesis for streptococcal and staphylococcal strains of similar *Lux*-cassette design. Furthermore, we challenged these microbes with different stimuli (*i.e*., myoinositol, oxamate, and carbon sources). We monitored their relative light production to dissect processes that would promote or repress the light production pathway in these strains. Given our results, it may be possible to use these light-producing pathogens as biosensors for the rapid assessment of physiologic processes and the local microbial environment.

## Results

### *D*-Glucose inhibits bioluminescence from *Strep.* Xen20

Our previous results indicated that the luminescence of *Strep.* Xen20 did not accurately reflect bacterial load *in vivo* and suggested this may be due to a *D*-glucose-mediated inhibitory effect (4). To confirm this finding on a static medium, we plated *Strep*. Xen20 on Todd-Hewitt with yeast extract (THY) plates supplemented with 0-or 50-mM *D*-glucose. For this experiment, cells were grown overnight and imaged for luminescence (Figure 1). Colonies with *D*-glucose supplementation showed decreased luminescence. Interestingly, the presence of sheep blood (SB) also significantly reduced the bioluminescence produced by *Strep.* Xen20. Similar CFUs were also observed by colony counting.

**Figure 1.**
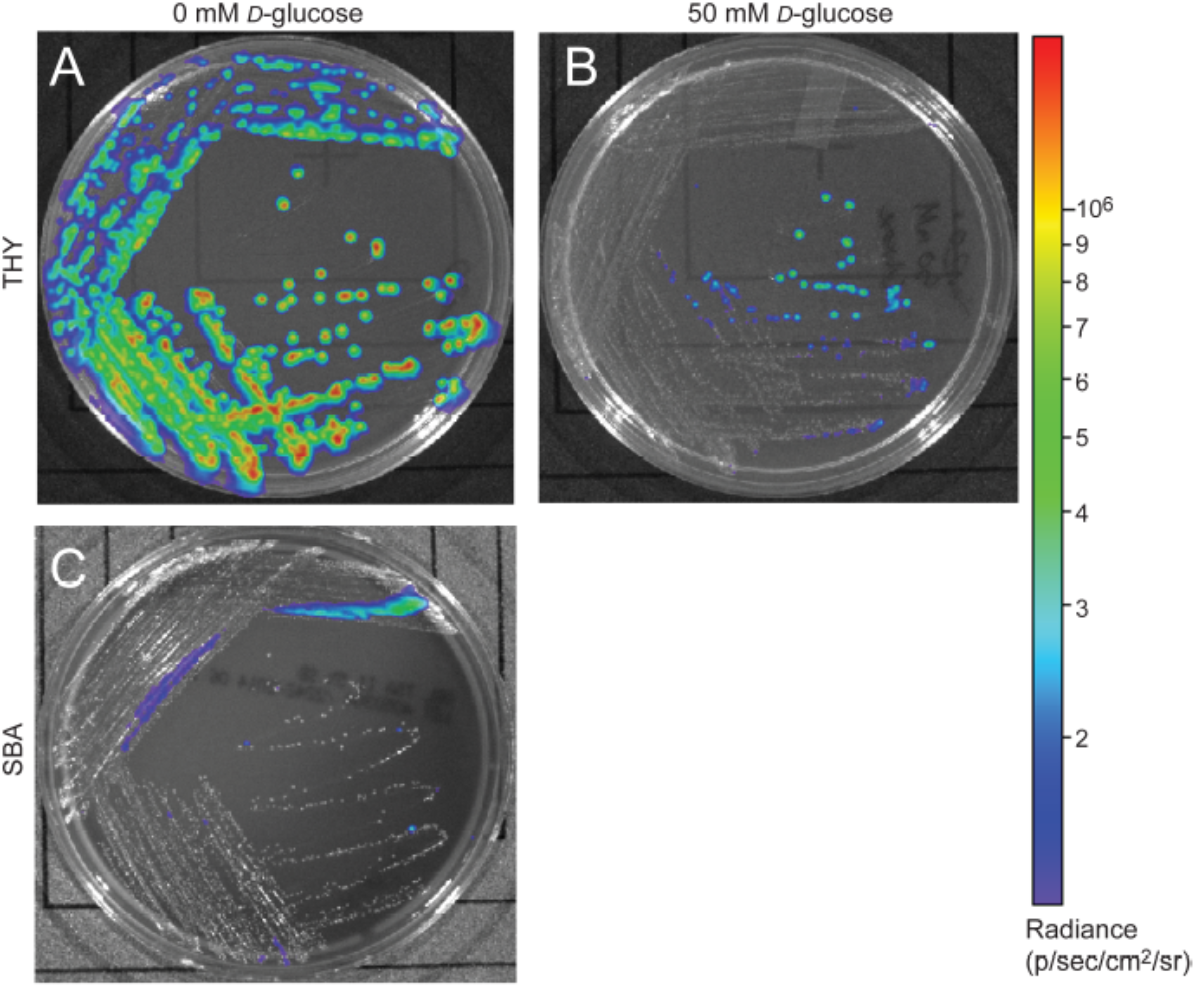
Glucose effect on bioluminescent production of *S. pyogenes* Xen20. *Strep.* Xen20 was plated on Todd Hewitt with yeast extract (THY) without (Panel A) or with the addition of excess D-glucose (Panel B) or on sheep blood agar (SBA) and incubated for 20 hours at 37 C.

### The lux operon in Xen20 is regulated by inositol and glucose levels

*Strep.* Xen20 has the *luxABCDE* cassette inserted in the *iol* operon used for inositol catabolism. More specifically, the gene is under the *iolE* promoter (8), which encodes 2-keto-*myo*-inositol dehydratase (9). The IolR repressor controls the entire *iol* operon (10). Under high inositol conditions, the repressor is liberated, and genes are expressed. As such, we tested the contribution of inositol to bioluminescence expression in *Strep*. Xen20. The results (Figure 2A) indicate increased light from bacterial cells grown in the presence of *myo*-inositol. In contrast, co-supplementation of *Strep.* Xen 20, with both inositol and a low concentration of glucose (3 mM), overcame this inositol-dependent enhancement. All BLI signal returned to baseline in less than 3 hours (Figure 2B). To evaluate this inositol benefit in actively growing cells, equal portions of *Strep*. Xen 20 were spread on Luria broth (LB) plates in the absence or the presence of 6mM myo-Inositol, of 50 mM *D-g*lucose, or both additives (Figure 2C). Results paralleled Figure 1 and the tube tests in Figure 2A-B. BLI from bacterial plates at 14, 24, and 38 hours in the presence of *D-*glucose plates showed near-complete loss of light production, and this was only slightly counteracted by the additional of myo-inositol at the 38 hour time point (Figure 2C).

**Figure 2.**
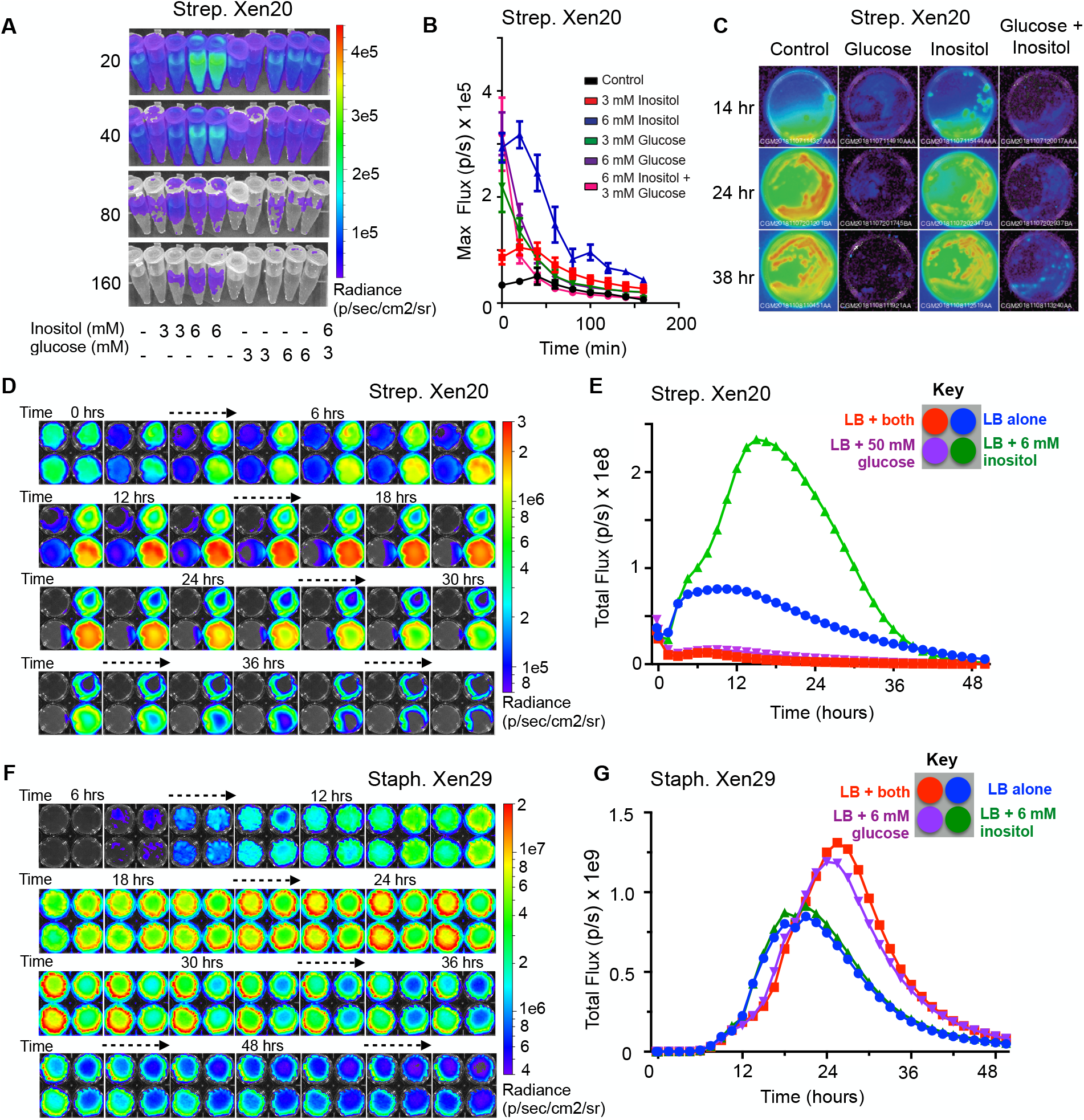
*Myo*-inositol regulates luminescence of *S. pyogenes* Xen20. *A.* Representative time course of *Strep.* Xen20 cells. Samples were grown to stationary phase and then diluted in fresh PBS supplemented with one of either 3 or 6mM *myo*-inositol or 3 or 6mM *D*-glucose. *B.* Quantification of luminescent signals from samples shown in Panel A shown as the maximum rate of photons generated (*Max flux*). *C.* BLI of 65mm LB plates containing no additive (*control*), 50mM glucose, 6mM myo-inositol, or both taken at 14, 24, and 38 hours after incubating at 37 C. *D. Strep.* Xen20 growth on LB plates containing either no additive (*control*), 50mM glucose, 6mM myo-Inositol, or both and imaged over 48 hours in the IVIS. *E.* BLI quantification of the plates from *Panel D* as the total rate of photons generates *(Total flux) F. Staph.* Xen29 incubated on LB plates containing either no additive (*control*), 50mM glucose, 6mM myo-Inositol, or both and imaged over 60 hours in the IVIS. *G.* BLI quantification of the plates from *Panel D* as the total rate of photons generates *(Total flux)*.

To support cross-conditional comparisons and to coordinate timing, we imaged the four distinct conditions simultaneously over two days—the *Strep*. Xen 20 kinetic profile showed a definite 2-reaction trace with the development of a new inositol peak at 16-17 hours that was almost ~4-fold greater than the amount of light emitted by the LB control (Figure 2D-E). Despite similar bacterial growth over the time course, the BLI signal for both the 50mM *D-*glucose and 6mM myo-inositol supplemented conditions showed markedly lower light production. Conversely, *Staph.* Xen 29 showed no change in light production with 6mM inositol supplementation alone and an ~2-fold increase when grown in the presence of *D-*glucose that was inositol-independent (Figure 2F-G). These results agree with previous research that showed the expression of green fluorescent protein (GFP), placed under the *iolE* promoter, was decreased in glucose-rich conditions under the *iolE* promoter in *Salmonella enterica* serovar Typhimurium strain 14028 (11).

### Substrate sources for bioluminescence of Streptococcus pyogenes

Since inositol cannot function as a carbon source in *Streptococcus pyogenes* M49 strains (based on the Kypto Encyclopedia of Genes and Genomes (KEGG) genomic data), we investigated the agonistic or antagonistic effects of central metabolites to the production of light. As previously stated, bacterial bioluminescence is dependent on the production of substrates by the bacterial cell (6); therefore, we investigated the link of bioluminescence to central metabolism in *S. pyogenes* (see pathway scheme in Figure 3). Bioluminescence is entirely dependent on the availability of activated acyl donors, which are produced *de novo* by the bacterial cell and serve as the substrate for the *lux* operon to create fatty aldehydes. These substrates are then converted to fatty acids in the presence of FMNH2 and oxygen, thereby releasing light. Activated acyl donors are synthesized in one of two ways: first, coenzyme A (CoA)-containing acyl compounds can be created via the breakdown of fatty acids by *β*-oxidation (green box, Figure 3); second, CoA-containing fatty acid building blocks can be created via the fatty acid biosynthesis pathway (salmon box, Figure 3). The second pathway is dependent upon the creation of acetyl-CoA by the glycolysis/lactic acid fermentation pathway (blue box, Figure 3).

**Figure 3.**
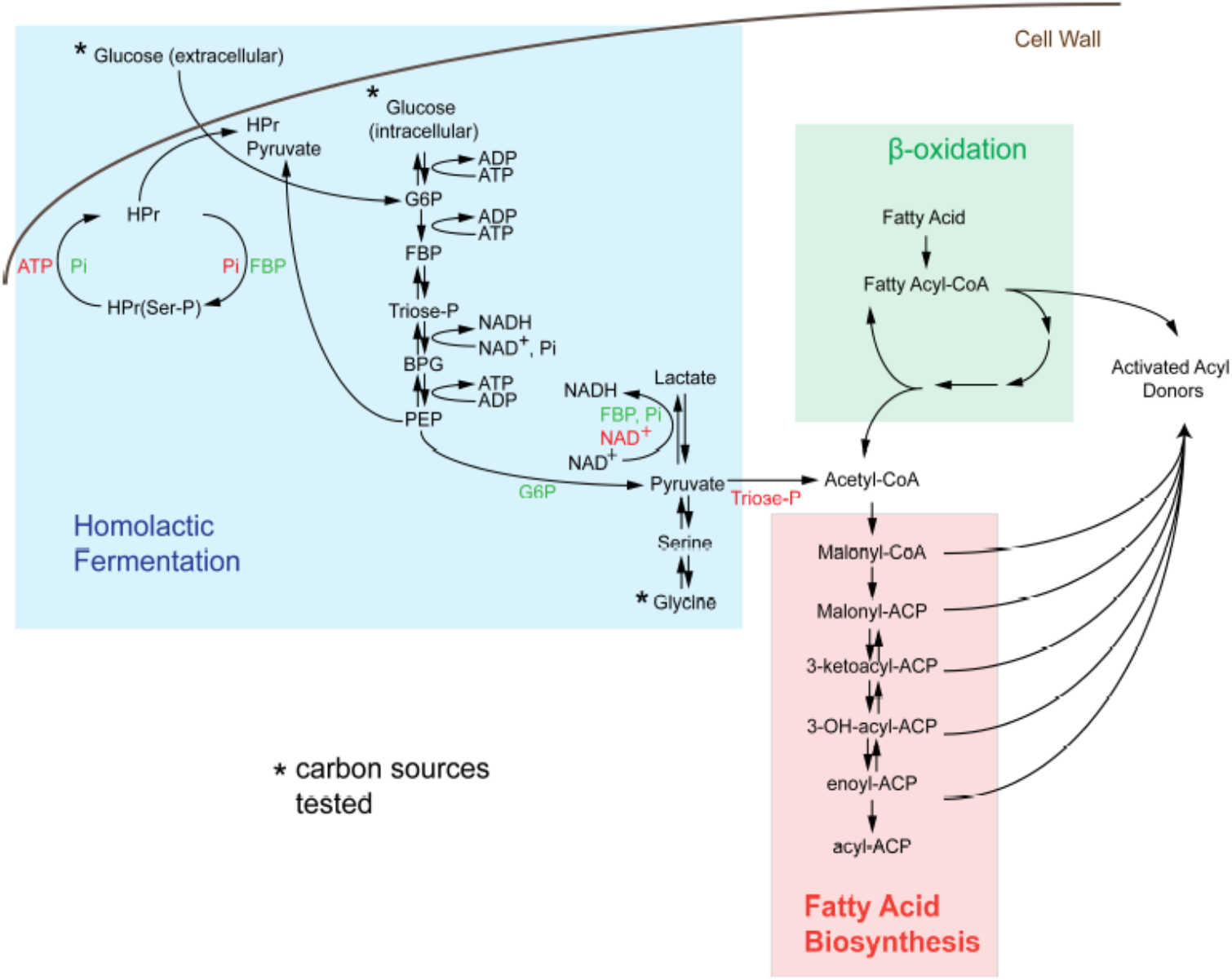
*De novo* synthesis of acyl donors via central carbon metabolism in *Streptococcus*. Activated acyl donors, the substrates for the *luxABCDE* machinery, are made by the breakdown of fatty acids through *β*-oxidation (*green region*) or by the synthesis of an activated carrier protein (*ACP*)-containing metabolites via fatty acid biosynthesis (*salmon region*). Since *S. pyogenes* lacks the *β*-oxidation pathway, all substrates for light production in *Strep*. Xen20 are created by acetyl-coenzyme A (*CoA*)-dependent fatty acid biosynthesis generated by homolactic fermentation (*blue box*). The asterisk indicates the sole carbon sources tested. Allosteric activators, such as inorganic phosphate (*Pi*), fructose bisphosphate (*FBP*), and glucose-6-phosphate (*G6P*) are shown in green text, and allosteric inhibitors, namely adenosine triphosphate (*ATP*), nicotinamide adenine dinucleotide (*NAD^+^*), and triose phosphate (*Triose-P*) are shown in *red text*. Abbreviation used in here is as following: histidine-containing carrier protein (*HPr*), phosphorylated HPr (*HPr(Ser-P*)), phosphoenolpyruvate (*PEP*), and 1,3-bisphosphoglyceric acid (*BPG*).

Although the KEGG Pathways are not available for *Strep.* Xen20, a pathway exists for the closely related M49 serotype ancestor NZ131 (http://www.genome.jp/dbget-bin/www_bget?genome:T00780) (12). Analysis of this pathway revealed important allosteric and feedback mechanisms for bioluminescence production. First, M49 strains lack the *β*-oxidation path beyond the creation of hexadecanoyl-CoA from hexadentate. In contrast, *S. aureus* NCTC8325, the parental strain of a frequently manipulated strain RN4220, showed increased capacity for degradation of fatty acids. Therefore, in M49 strains, the majority of activated acyl donors must be created by the fatty acid biosynthesis pathway. Analysis of these pathways revealed M49 to have all necessary enzymes for these pathways (Supplemental Figure 2), which led us to investigate the contribution of glycolytic compounds on the BLI signal.

### Dependence of luminescence on glucose homeostasis

*Strep.* Xen20 processes sugars by homolactic fermentation, a process that utilizes *D*-glucose as the preferred carbon source (13). Based on KEGG genome available data, glycine is converted to pyruvate via conversion to serine by L-serine dehydratase in reactions essential to the formation of one-carbon metabolites.

We, therefore, utilized *D-*glucose as a glycolytic carbon source and glycine as a non-glycolytic carbon source. *D*-Glucose is present at physiological concentrations ranging between 3.9-5 mM in healthy adult mice (14). Therefore, we tested exogenous *D*-glucose and glycine at either 3 or 6 mM (Figure 4A). The addition of either exogenous *D*-glucose and glycine to *Strep.* Xen20 decreased the BLI signal. Counter to this, the addition of *D*-glucose or glycine increased luminescence expression in *Staph.* Xen29 (Figure 4B).

**Figure 4.**
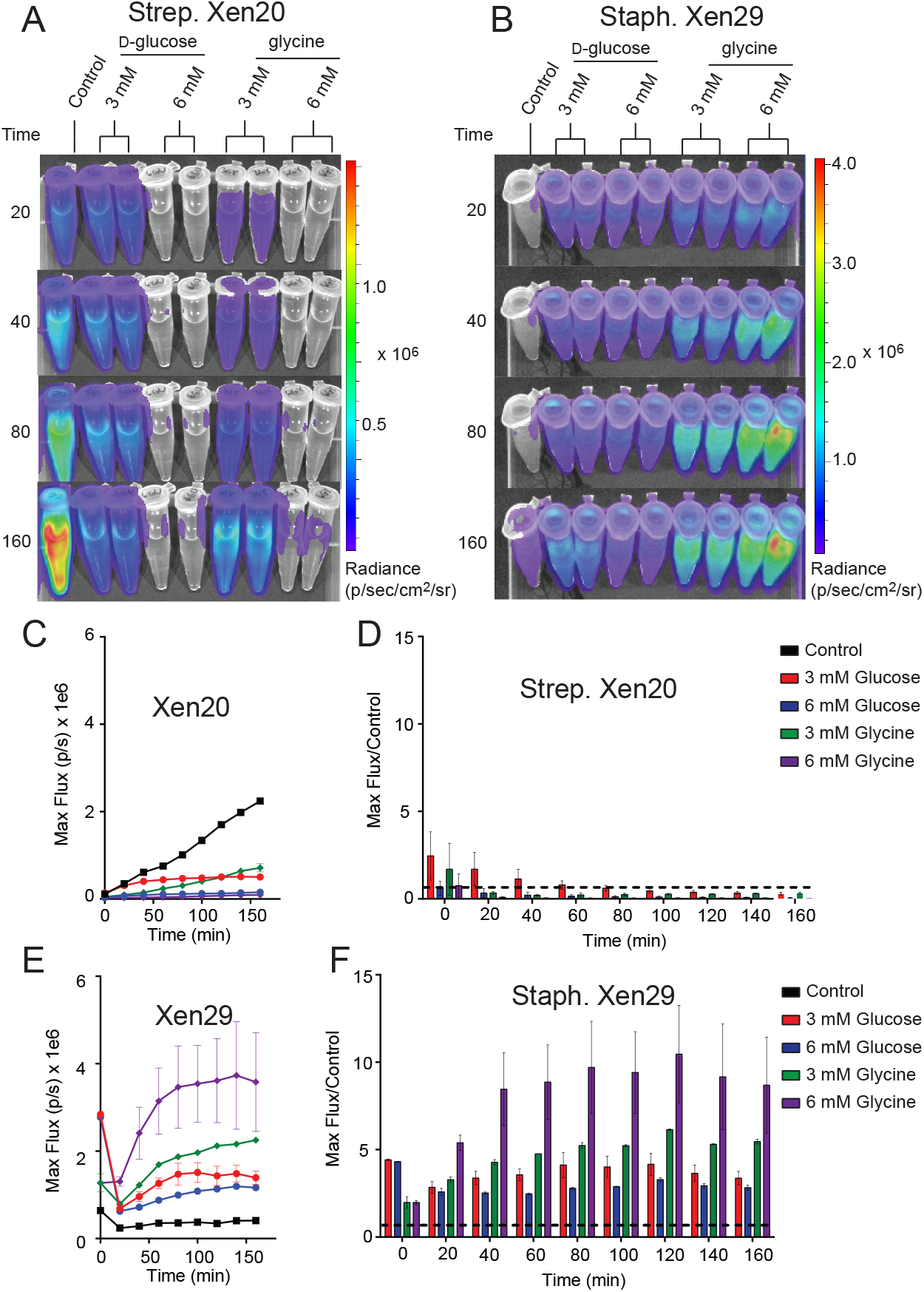
The effect of monosaccharides on bioluminescence production by Gram-positive pathogen. *A*. Representative time course of *Strep*. Xen20 or *Staph.* Xen29. For all samples, cells were grown to stationary phase then diluted in fresh phosphate-buffered saline (*Control*) supplemented with either *D*-glucose or glycine as indicated. *B*. Similar time course for Xen29 (*S. aureus*). *C*. Quantification of the *Strep*. Xen20 luminescent signal from Panel A. *C-D*. Quantification of luminescent signal from samples shown in Panel A for both *Strep*. Xen20 and *Staph.* Xen29 pathogens represented as either the maximum rate of photons generated (shown on the *left* as *Max flux*) or as the relative maximum signal normalized to the control (shown on the *right* as *Flux/Control*) displayed as a function of time. The black dashed line indicates a ratio at the threshold of 1.0. *E-F*. Similar quantification of Panel B for Staph. Xen29. All samples are color coded as indicated and the experiment performed, as shown in the Materials and Methods section.

### Physiologically relevant limitations in bioluminescence can be attributed solely to metabolites upstream of pyruvate

To test tissue-derived carbon sources, we perturbed the carbon source equilibrium of both light-producing pathogens (*Strep.* Xen20 and *Staph.* Xen29) by characterizing the glucose-dependency of BLI signal. The dose-dependent effect of *D*-glucose on light production was monitored following incubation with M9 minimal media supplemented only with casein hydrolysate and yeast extract (Figure 5). The supplements meant to provide the microbe with amino acids and cofactors essential for protein and DNA synthesis. In this moderately enhanced M9 medium, we found that Xen20 produced detectable levels of luminescence. Providing an additional glycolytic carbon source, in the form of *D-*glucose, did not significantly increase radiance. Contrastingly, Xen29 produced only moderate levels of luminescence in the absence of further carbon sources. The addition of 6 mM *D-*glucose significantly increased the production of light in Xen29. Similar to *D-* glucose, Xen20 grown in modified M9 alternatively supplemented with glycine (3 mM) produced no significant increase in BLI signal. In contrast, all supplementations increased bioluminescent expression in *Staph. Xen29* cells with maximal BLI signal when glycine was present (Figure 5).

**Figure 5.**
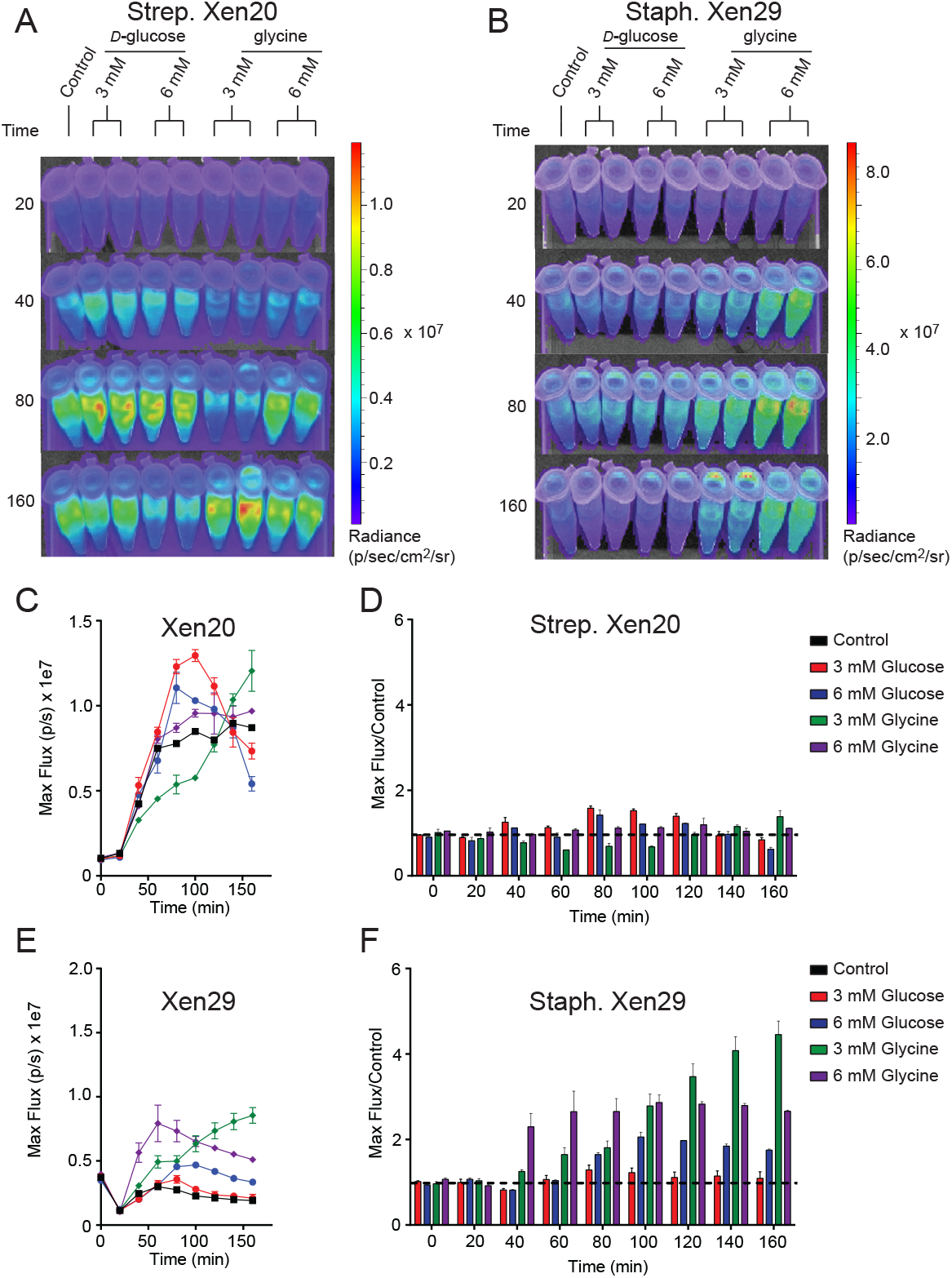
Increased luminescence production by Gram-positive pathogen incubated in M9 medium supplemented with casein hydrolysate and yeast extract. OD_600nm_ and luminescence were measured using a Thermo Fischer VarioSkan plate at 20-minute intervals over 160 minutes at a constant 37 °C. *A*. Representative time course of *Strep.* Xen20. For all samples, cells were grown to stationary phase then diluted in fresh M9 medium containing casein hydrolysate and yeast extract (*control*) supplemented with either *D*-glucose or glycine as indicated. *B*. Similar time course for Xen29 (*S. aureus*). *C*. Quantification of the *Strep*. Xen20 luminescent signal from Panel A. *C-D*. Quantification of the *Strep*. Xen20 luminescent signal from Panel A represented as either the maximum rate of photons per second (*Max flux*) and as the relative maximum signal normalized to the control (shown on the *right* as *Flux/Control*) displayed as a function of time. The black dashed line indicates a ratio at the threshold of 1.0. *E-F*. Similar quantification of Panel B for Staph. Xen29. All samples are color coded as indicated and the experiment performed, as shown in the Materials and Methods section.

### Restriction of luminescence is dependent on L-lactate dehydrogenase

Homolactic fermentation produces pyruvate that is either converted to acetyl-CoA by pyruvate dehydrogenase or to lactate by L-lactate dehydrogenase (*blue box*, Figure 3). Pyruvate conversion by pyruvate dehydrogenase would prepare substrates relevant to light production. In contrast, conversion by L-lactate dehydrogenase would scavenge pyruvate away from this potential pool. To test the effect of pyruvate conversion to lactate and the availability of the acetyl-CoA pool, various concentrations (0-64 mM) of oxamate, an inhibitor of L-lactate dehydrogenase, were added to *Strep*. Xen20 cells (Figure 6) (15). However, luminescence was constant up to 32 mM oxamate (Figure 6). At the 64mM oxamate, the radiance increased for the glucose-containing sample, but there was also an observed decrease in cell density. A diffusion assay tested the effect that drug eluted in to agar has on light production (Figure 6B). Equal CFUs of *Strep.* Xen20 cells were added to the quadrants of an agar plate, and an oxamate or relevant control discs were added (Figure 6B). Discs were then imbued with equivalent volumes applied to them (20μl), and the solutions were sterile water, glucose alone (0.022 mg), oxamate (0.14 mg), and both glucose and oxamate. Dried discs were placed on the agar plates. Bioluminescence production was higher in quadrants with oxamate (Figure 6C).

**Figure 6.**
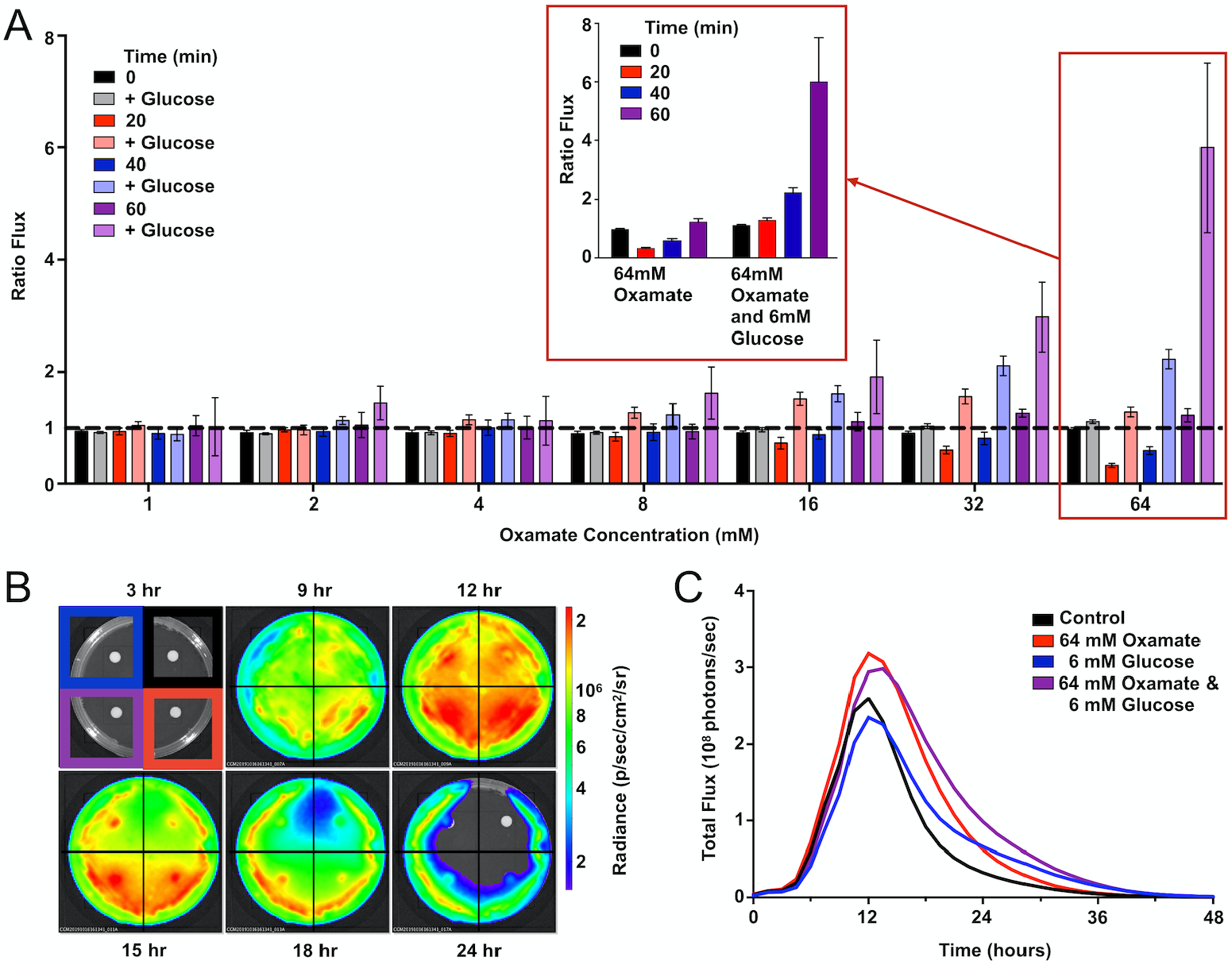
Oxamate inhibition partially recovers the bioluminescent signal. *A. Strep*. Xen20 in the stationary phase were combined with PBS alone *(control)* or PBS with concentrations of 6 mM glucose and 0-64 mM oxamate. Represented is the flux from cells treated with oxamate divided by the flux from cells in PBS alone *(Ratio Flux)*. The data represents seven trials without or without glucose at the indicated times. Expansion of the 64 mM oxamate concentration with and without glucose over 60 minutes shown *(red inset)*. *B. Strep*. Xen20 was grown on LB agar with disks containing either a water control *(upper right)*, 64 mM oxamate (*lower right)*, 6 mM glucose *(upper left)*, or both glucose and oxamate *(lower left)* monitored for 48 hours. *C.* Quantification of the luminescent signal from Panel C as the total rate of photons generates *(Total flux)*.

### Determining plasma glucose concentrations based on relative luminescence

Given the dependence of Xen20 bioluminescence on glucose homeostasis, we sought to correlate glucose concentration with the relative light production of *Strep.* Xen20. Light production of the Xen20 decreased at relatively similar rates when added to an equal volume of glucose in phosphate buffered saline (PBS). However, each sample displayed a different time to initial decrease and terminal luminescence (Figure 7A). Glucose concentration in the PBS and mouse plasma samples was confirmed using a commercial blood glucose meter. Based on the linear relationship, the mouse plasma had a glucose concentration of 3.3mM (Figure 7B). After the D-glucose standards were allowed to incubate with the *Strep.* Xen20 culture overnight, a exponentially decreasing relationship was found to exist between the amount of glucose and *Strep.* Xen20 BLI (Figure 7C). Data was analyzed using a single exponential and used to predict glucose concentration in mouse plasma. Our predicted concentration of glucose in the mouse plasma matched actual glucometer values determined independently.

**Figure 7.**
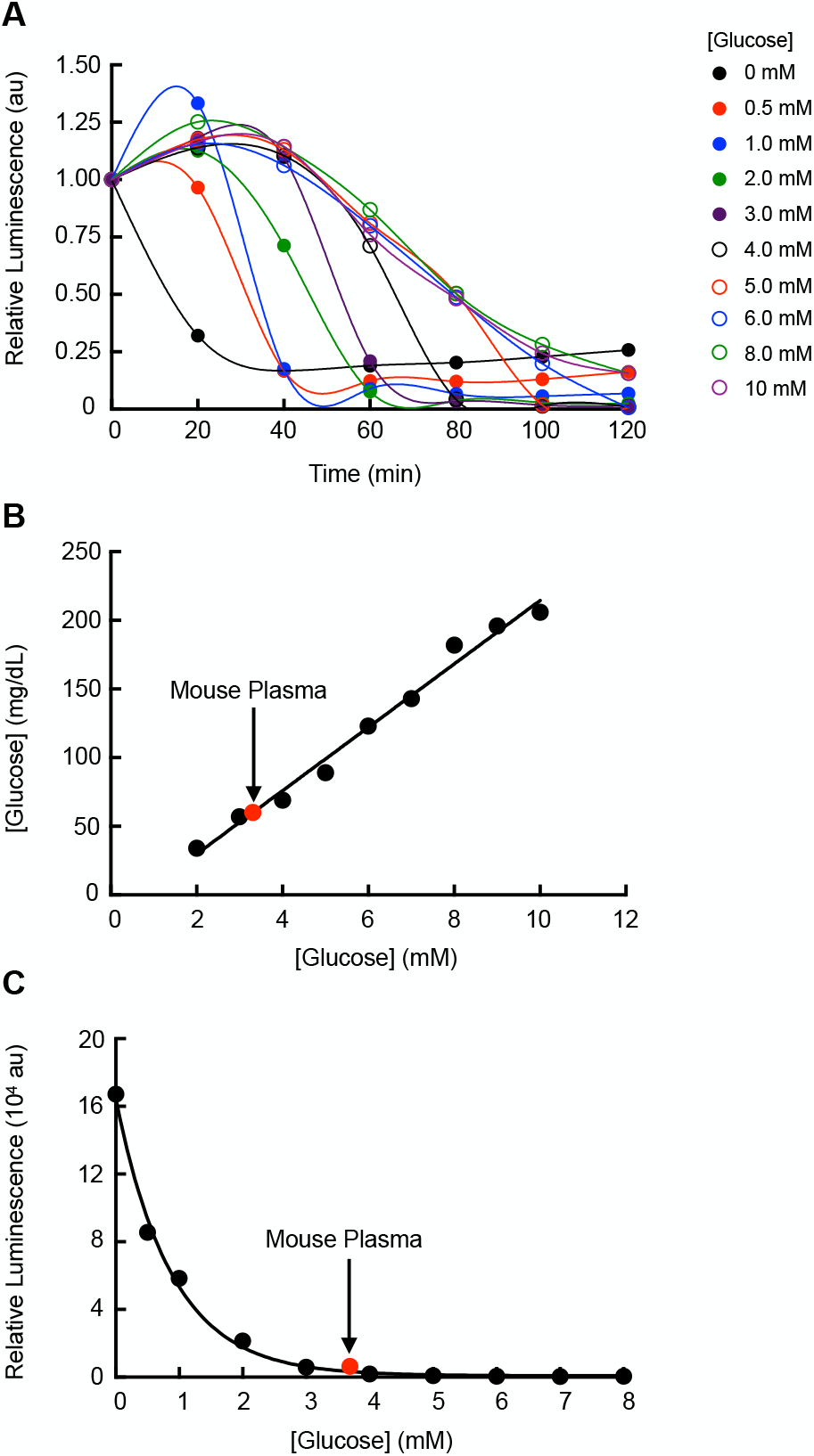
Utilizing *Strep.* Xen20 bioluminescence production to estimate plasma glucose concentrations. *A. Strep.* Xen20 cells in the stationary phase were added to equal volumes of PBS with 0-10 mM glucose. The luminescence was measured with a luminometer every 20 minutes for 2 hours. *B.* The glucoses level (mg/dL) of each PBS and glucose solution was measured using a commercially available blood glucose meter. Mouse plasma (indicated on graph in red) was also tested for blood glucose levels, which was used to estimate millimolar concentration. *C.* Final luminescence in the *Strep.* Xen20 sample in glucose between 0 and 8mM after incubation overnight. The exponential relationship yielded an equation of y=167000e^−0.882x^ (R^2^=0.9861) and was used to estimate the mouse plasma glucose level based on luminescence of the *Strep.* Xen20.

## Discussion

Competition for carbon sources and energy may affect the survival of *Streptococcus pyogenes* (*S. pyogenes*) during an infection. Previously we found that a *luxABCDE*-incorporated *S. pyogenes* had reduced light production in *D*-glucose enriched medium (4). Although this trait is less desirable for *Strep.* Xen20 utility in non-invasive imaging of infections, it highlights a unique ability of bacterial luciferase to serve as a biosensor of activated or repressed metabolic pathways. We explore the use of *Strep.* Xen 20 as a biosensor that could respond to changes in its local environment by altering its light production. This biosensor would fundamentally differ from the current utilization of isolated luciferases in biochemical assays. Specifically, our biosensor reports the activity of metabolic enzymes (substrates for light production) rather than on the level of promoter activity intrinsic to firefly or Renilla luciferase (under a bolus of the substrate).

Initially, genetic evidence pointed to *iolE* as the gene sequence interrupted by the *luxABCDE* pathway (8). Results of inositol supplementation (Figure 2) indicate a genetic regulation of the *luxABCDE* bioluminescence. According to the KEGG metabolic pathways for NZ131, M49 serotype strains of *Strep.* Xen20 does not contain the enzymes necessary to convert this inositol further into usable carbon sources to enter glycolysis or citric acid cycle. Therefore, we tested the effects of other carbon sources to link the creation of luminescent substrates to the central metabolism of the bacterial cell.

Our analysis of the KEGG metabolic pathways for NZ131 revealed no enzymes capable of fatty acid degradation beyond the creation of hexadecanoyl-coenzyme A (see the scheme in Figure 3). Necessary activated fatty acyl compounds must then be created via the fatty acid biosynthesis pathway, which utilizes acetyl-CoA as building blocks for the creation of fatty acid chains. As previously mentioned, acetyl-CoA is produced in *S. pyogenes* via homolactic fermentation. Taken together, this suggests *S. pyogenes* has a more simplified route to the production of the essential light-producing building block than its counterpart, *S. aureus*.

Glucose is utilized in one of two ways: first, it can be transported into the cell by phosphotransferases, which convert it to glucose-6-phosphate during uptake, and fed into the fermentation process; second, it can be converted to glucose-6-phosphate by hexokinase (13). The addition of *D*-glucose and glycine were shown here to decrease the bioluminescence of *Strep.* Xen20. As seen in Figure 3, and reviewed elsewhere (13), excess glucose during homolactic fermentation causes the conversion of pyruvate to lactate by the enzyme L-lactate dehydrogenase (LDH) due to increased levels of fructose-bisphosphate (FBP), intensified by the inhibition of pyruvate conversion to acetyl-CoA by increased triose phosphate levels. Therefore, the decreased bioluminescence observed in *D*-glucose supplementation may be due to the reduced acetyl-CoA pool, and subsequent lack of fatty acid biosynthesis precursors. As such, we tested the addition of an inhibitor of LDH to attempt to restore the acetyl-CoA pool and improve the luminescent signal (15). Oxamate-containing discs allowed for the increased light production in the dispersion area (Figure 6). Therefore, bioluminescence is diminished in the presence of increased glucose due at least in part to its generation of FBP. This FBP acts on LDH to scavenge the pool of available pyruvate, committing it to the production of lactate rather than the fatty acyl precursor acetyl-CoA. Glycine supplementation showed comparable bioluminescence levels at concentrations equal to those of *D*-glucose. Although glycine is first converted to pyruvate, gluconeogenesis can convert this pyruvate back to triose phosphate and/or FBP, creating a similar effect to *D*-glucose supplementation.

Interestingly, the addition of amino acid sources, such as casein hydrolysate, relaxed the inhibition of bioluminescent signal by *D*-glucose and glycine supplementation (Figure 5). According to the KEGG diagram for *S. pyogenes* NZ131, L-alanine can be converted to pyruvate and contribute to luminescence. In contrast, amino acids that are converted to fumarate are unable to contribute due to the lack of citrate cycle activity. Leucine, valine, and isoleucine are converted to their subsequent oxopentanoates, but subsequent reactions are not possible due to a lack of 3-methyl-2-oxobutanoate dehydrogenase. It is unclear which amino acid or a combination of amino acids plays a role in increasing the *Strep.* Xen20 light production in the presence of glucose and, as such, dissection of that would require further study.

The application of the data gathered here led us to investigate how *Strep.* Xen20 might be used as a living sensor of local glucose concentration. The results of our glucose standards and mouse plasma glucose levels give proof-of-concept evidence that *Strep.* Xen20 can be used to determine the approximate levels of blood glucose in a given sample, similar to the heme sensing capabilities of the luminescent *E. coli*. (5) Here, we evaluated the *Strep.* Xen20 luminescence using mouse plasma. However, it is entirely feasible that *Strep.* Xen20 could detect glucose in other species and in the host.

To our knowledge, these results are the first to highlight the effects of the central metabolism on the bacterial luciferase system. It is conceivable that directed insertion of the luxABCDE operon could generate a biosensor that reporters on the presence of trace heavy metals or the presence of toxicity compounds (such as arsenic). It would be easy to ignore the complications of light production by using non-integrated plasmid versions of LuxAB in these pathogens and injection luciferin substrate, such as the *ffluc* system previously described in *S. pyogenes* (16). This would be simpler, in many regards, but future correlations of metabolism-dependent light production with RNA-seq technology could provide new avenues for the high-throughput assessment of compounds that disrupt molecular pathways regulating these dangerous pathogens. These would help identify non-redundant processes in these bacteria that if targets would not adversely affect human/host metabolism.

## Experimental Procedures

### Chemicals and Reagents

Brain-Heart Infusion (BHI), Sheep Blood Agar (SBA), and Todd-Hewitt with yeast extract (THY) were from BD Biosciences (San Jose, CA) or RPI. Kanamycin and *D*-glucose were purchased from Research Products International (RPI; Mt. Prospect, IL). Glycine was purchased from AMERSCO (Solon, OH). Unless otherwise specified, all reagents were purchased from Sigma Aldrich. M9 minimal medium was prepared as previously described (17).

### Bacterial Strains, Cultivation Conditions, and Imaging

Strains *Strep.* Xen20 and *Staph.* Xen29 (PerkinElmer Inc., Waltham, MA) were grown to stationary phase (OD_600nm_ > 1) in BHI broth for 18 hours at 37 °C and approximate concentration determined by light scattering at OD_600nm_ per manufacturer instructions. Cultivation conditions were altered either in the liquid media or on solid media and supplemented with additive, as indicated. Plates and tubes were imaged on the IVIS Lumina XRMS or Lumina II system (PerkinElmer Inc.). BLI is reported as calibrated units of radiance (p/sec/cm2/sr) in LivingImage software version 4.7 (PerkinElmer Inc.), allowing for comparisons between detectors.

### Bioluminescence Kinetic Assays

To test if the *lux* operon was indeed under the inositol promoter (*iolE*) (8), we diluted bacteria in PBS containing *myo*-inositol, *D-glucose,* or both in a final 1mL volume at equivalent OD_600nm_. Tubes were incubated and imaged in the IVIS Lumina XRMS on a 37 °C stage and imaged every 20-minutes for 160 minutes. *Strep*. Xen20 was also streaked on a secondary set of plates with Luria broth (LB, RPI) alone or supplemented with either 6 mM myo-inositol, 50 mM *D*-glucose, or both myo-inositol and *D*-glucose. Plates were incubated at 37°C and imaged at 14, 24, and 38 hours. Catabolite repression was assessed by supplementation of agar with myo-inositol or *D-*glucose or both in 6-well, flat-bottomed culture plates (Costar, Corning, NY) and imaged every 1.5 hours with the IVIS Lumina II for 48 hours.

The effect of carbon source availability was determined by incubating cells in sterile PBS (control) or PBS-supplemented with either 3 mM or 6 mM of *D*-glucose or glycine. Sample tubes were diluted and imaged as before. To determine the contribution of growth rate to *Strep*. Xen20 expression, this process was repeated for M9 minimal medium supplemented with 1% casein hydrolysate and 0.3% yeast extract. This media provides necessary cofactors for growth in the absence of confounding carbon sources.

### Oxamate Inhibition Assay

To determine the contribution of L-lactate dehydrogenase on *Strep.* Xen20 bioluminescence, *Strep*. Xen20 cells in the stationary phase were added to PBS containing a final concentration of 6 mM *D*-glucose in a 96-well plate. Varying amounts of oxamate were added for final concentrations of 0-256 mM. OD_600nm_ and luminescence were measured using a Thermo Fischer VarioSkan plate at 20-minute intervals over 160 minutes at a constant 37 °C. For comparison purposes, cells treated with oxamate were compared to those untreated in both the control and glucose-supplemented groups. In addition, *Strep.* Xen20 was grown and imaged for bioluminescence over 48 hours on an LB agar plate containing discs saturated with 20mcL of solutions containing 6 mM oxamate, 6 mM *D*-glucose, or both. The discs were prepared by the addition of 4 equal volume solutions (20μl) corresponding to sterile water, glucose alone (0.022 mg), oxamate (0.14 mg), and both glucose and oxamate. The discs were dried before placing them on the agar plates.

### Plasma Glucose Effect on Bioluminescence

To evaluate *Strep.* Xen20 as a potential monitor of local plasma glucose concentrations, *Strep.* Xen20 grown to the stationary phase in BHI was combined in a 1:1 ratio with glucose in PBS at concentrations between 0 and 10 mM. *Strep.* Xen20 was also combined with C57BL6 mouse plasma (Innovative Research, Novi, MI). The light production was then monitored at 20-minute intervals at 37 °C for 2 hours using the Glomax 20/20 Luminometer (Promega, Madison, WI). A final measurement was also taken after the samples had been allowed to incubate overnight at room temperature. The glucose level in the glucose standards and mouse plasma were also measured directly using a commercially available ReliOn PRIME blood glucose meter (Wal-Mart Stores, Inc. Bentonville, AR) ReliOn™ PRIME blood glucose test strips. The meter was unable to determine the glucose levels at < 2 mM.

### Statistical Analysis

Statistical means of each group were analyzed using two-way, repeated-measures analysis of variance (ANOVA) on R 3.1.2 (http://www.r-statistics.com/on/r/). Time, substrate, and the interaction of time*substrate were analyzed for each medium and each bacterium. Results were reported for statistical values p < 0.05, or no significant difference for p > 0.05.

### Data availability

All remaining data are contained within the article and Supporting Information

## Abbreviations

*S. pyogenes*: Streptococcus pyogenes
*luxABCDE*: *Photorhabdus luminescens lux* operon cassette
*ffluc*: firefly luciferase
*Strep*. Xen20: *Streptococcus pyogenes* Xen20;
*E. coli*: *Escherichia coli*
FMNH_2_: reduced flavin mononucleotide
THY: Todd-Hewitt with yeast extract
SB: sheep blood
GFP: green fluorescent protein
KEGG: Kypto Encyclopedia of Genes and Genomes
CoA: coenzyme A
BHI: Brain-Heart Infusion
SBA: Sheep Blood Agar (SBA)
ANOVA: analysis of variance
LB: Luria broth
PBS: phosphate buffer saline
NAD^+^: nicotinamide adenine dinucleotide
triose-P: triose phosphate
HPr: histidine-containing carrier protein
HPr: (HPr(Ser-P)
PEP: phosphoenolpyruvate
BPG: 1,3-bisphosphoglyceric acid

## Funding and additional information

This research was supported in part by NIH R01-HL114477 to PRP. The content of the paper is solely the responsibility of the authors and does not necessarily represent the official views of the National Institutes of Health.

